# Northward expansion of the barnacle *Fistulobalanus albicostatus* in Japan

**DOI:** 10.64898/2026.07.06.736396

**Authors:** Masami M. Tamechika, Adnan Shahdadi, Benny K. K. Chan

## Abstract

*Fistulobalanus albicostatus* Pilsbry, 1916 (Thoracica: Balanidae) is a tropical to temperate species distributed in the NW Pacific. The previously known northernmost record of this species in Japan was from Aomori Prefecture, at the northern end of Honshu Island, Japan. However, field surveys conducted in 2023 and 2026 confirmed the occurrence of *F. albicostatus* in Hakodate Bay, the southern end of Hokkaido, Japan, across the Tsugaru Strait, thereby extending the northern limit of its known distribution. A line transect survey conducted in May 2026 recorded seven living individuals within an area of 128 m. *F. albicostatus* was rare on the mid-high shores, accounting for only 2% of all barnacle individuals in a quadrat survey. The basal diameter of the living individuals ranged from 0.76 to 1.23 cm, and all individuals possessed ovaries. Based on characteristics of both morphological and COI gene, the specimens were identified as *F. albicostatus*, and belonged to the same haplotype of populations that are present in Honshu Island. The establishment of *F. albicostatus* in Hokkaido suggests an ongoing northward range shift of this warm-water species, with the potential for further expansion under continued ocean warming.

## Introduction

Global warming and climate change are having profound effects on the geographical distributions of organisms (Chen *et al*., 2011; Pinsky *et al*., 2020). In marine ecosystems, new northernmost distribution records are increasingly being reported from high-latitude regions where successful establishment was previously considered unlikely (Sorte *et al*., 2010; Lenoir *et al*., 2020). Furthermore, cold range edges have been shown to shift more rapidly in response to climate change than warm edges (Fredston-Hermann *et al*., 2020). Cold water regions, show more rapidly changes in species diversity than tropical regions. The Arctic region has warming responses four times more than the tropical counterpart (Rantanen *et al*., 2022). In the past ten years, the blue mussels *Mytilus edulis* from the Atlantic invaded the Svalbard waters in the Arctic (Kotwicki *et al*., 2021). Rising temperatures can facilitate shifts in such cold range edges by reducing overwintering barriers, and thereby increasing the survival and establishment of tropical and warm water species in higher-latitude regions (Figueira and Booth, 2010). Consequently, accumulating knowledge of such range expansions is essential for understanding the geographic distributions of species and predicting future distributional shifts under continued climate warming.

*Fistulobalanus albicostatus* Pilsbry, 1916, is a tropical to warm-temperate barnacle widely distributed in East Asia, including Taiwan, southern China, the Korean Peninsula, and Japan (Pilsbry, 1916; Chang *et al*., 2017). This species is primarily found on mangrove trunks and leaves, as well as boulders and artificial structures in estuarine intertidal habitats (Utinomi, 1967; Chang and Leung, 2007). It has also been reported from cultured oysters and oil platforms (Pilsbry, 1916; Henry and McLaughlin, 1975; Foster and Willan, 1979). In addition, *Fistulobalanus* species have been documented attached to mobile animals such as migratory birds and sea turtles, which may facilitate long-distance dispersal (Tøttrup *et al*., 2010; Hayashi *et al*., 2017). As a warm-water species with multiple potential dispersal pathways, *F. albicostatus* may be undergoing expansion of its cold range edge in response to rising seawater temperatures.

The northernmost record of *F. albicostatus* is Mutsu Bay, Aomori Prefecture, Honshu Island, northern Japan (Utinomi, 1967). Likewise, a comprehensive survey of barnacle fauna in northern Japan, including Hokkaido, also recognized Mutsu Bay as its northern limit (Kado, 2003). *F. albicostatus* has not been documented from Hokkaido, beyond the Tsugaru Strait separating Honshu and Hokkaido (see Kado, 2003). However, we confirmed the presence of *F. albicostatus* in Hakodate Bay, Hokkaido (Figure 1A), in both 2023 and 2026. Here, we report the first northernmost record of the species from Hokkaido and extend its known northern distribution limit, quantifying its abundance using line-transect and quadrat surveys. In addition to our morphological identification, we utilized mitochondrial cytochrome c oxidase subunit I (COI) gene sequences to determine the phylogenetic affinity of the Hakodate population to previously identified subclades.

**Fig. 1.**
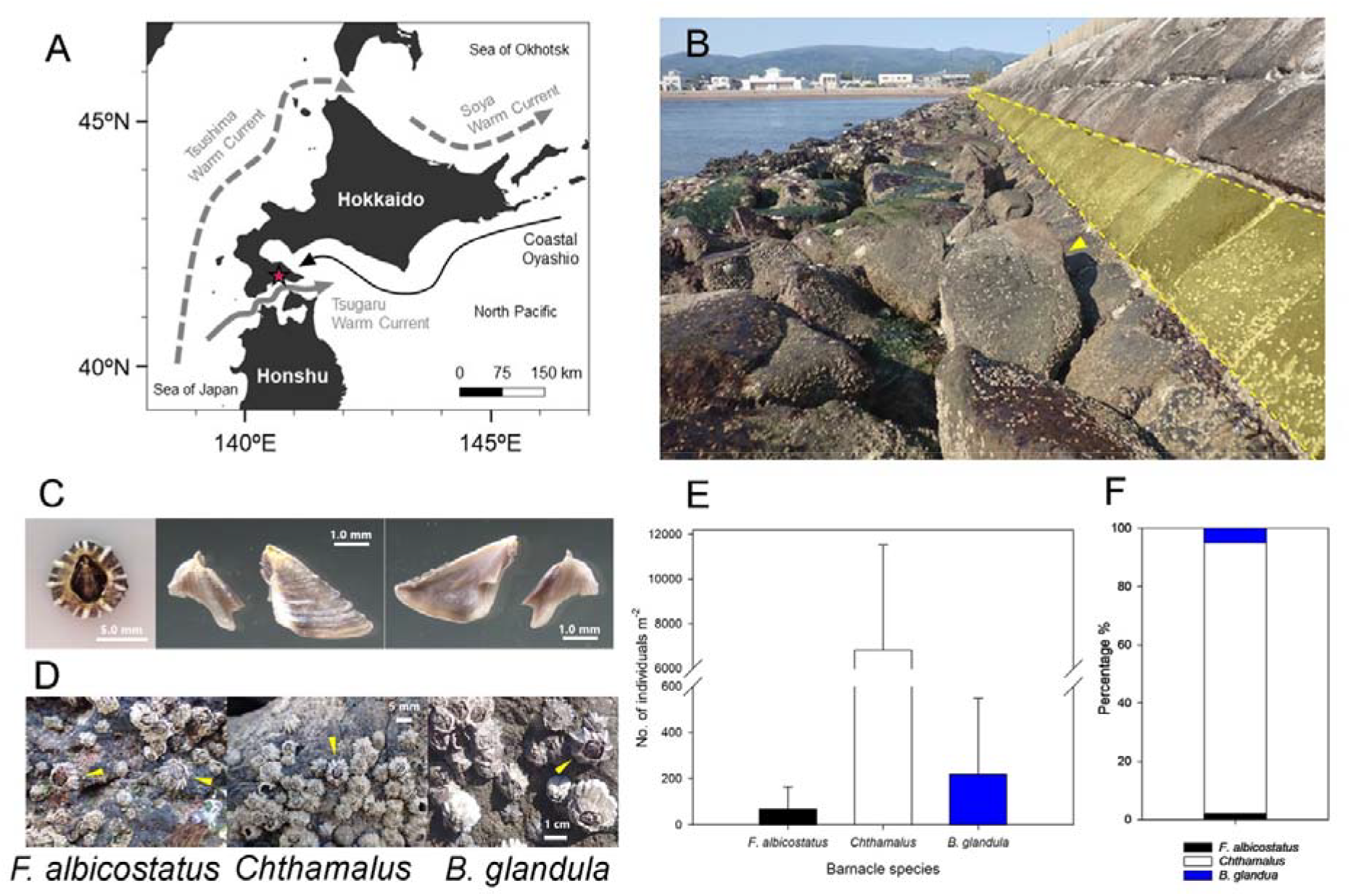
(A) Map of the area around Hokkaido, Japan, showing the major ocean currents and the study site (red star) in Hakodate Bay. Arrows indicate the main currents around Hokkaido. (B) The study site composed of an inclined seawall. Yellow shading indicates the line transect survey areas, and a yellow triangle indicates the location of an individual that was occasionally found on a boulder. Both the transect areas and the boulders were located at the tidal level where *Fistulobalanus albicostatus, Chthamalus* and *Balanus glandula* were found. (C) Left - External shell of *F. albicostatus*. Note the thick white rib, a diagnostic character for *F. albicostatus*. Middle - external view of tergum and scutum, right - internal view of scutum and tergum. (D) Close up view of *F. albicostatus*, with *Chthamalus* and *B. glandula*. Yellow triangles indicate the locations of *F. albicostatus* individuals. (E) Mean abundance of *F. albicostatus, Chthamalus* (including *C. dalli* and *C. challengeri*), and *B. glandula* from the transect survey at the study site. (F). Percentage of average abundance of *F. albicostatus, Chthamalus*, and *B. glandula* from the transect survey, showing *F. albicostatus* is a minority species in the intertidal zone.

## Materials and Methods

### Study sites and long-term temporal variations in seawater and air temperature

Field surveys were conducted at Nanaehama, Hakodate Bay, Hokkaido (41°48′43.6″N, 140°42′13.0″E; Figure 1A) on 23 May 2023, 28 December 2023, 22 May 2026, and 29 June 2026, within the intertidal zone, including both rocky shore and inclined seawall habitats.

All collected *F. albicostatus* individuals were preserved in 99.5% ethanol for subsequent COI sequencing. And then, a subset of specimens collected in May 2023 (*N* = 1) and May 2026 (*N* = 8) were measured for shell dimensions (aperture length, basal diameter, and shell height); the presence of ovaries was examined, and penis length measurements were conducted only for specimens collected in May 2026.

To examine long-term changes in climatic conditions in southern Hokkaido, surface seawater and air temperature from January 2000 to December 2025 were extracted from the database of Usujiri Fisheries Station, Hokkaido University. This temperature dataset was measured daily at 9:00 a.m. at Usujiri, southern Hokkaido (41°56′11.7″N, 140°56′59.9″E). Annual mean, maximum, and minimum surface seawater temperatures, and the annual number of days with surface seawater temperatures exceeding 20°C or below 5°C, were calculated to assess long-term changes in local climatic conditions. The same metrics were also calculated for air temperature using thresholds of 25°C and 0°C. Temporal trends in these climate conditions were examined in R version 4.4.2 (R Core Team, 2024). Annual mean, maximum, and minimum temperatures were analyzed using linear models (LMs). Annual number of days above and below the threshold temperatures were analyzed using generalized linear models (GLMs) with a negative binomial distribution and log-link implemented with the “glmmTMB” package (Brooks *et al*., 2017; McGillycuddy *et al*., 2025). These visualizations and their associated 95% confidence intervals were created using the “ggplot2” package (Hadley, 2016).

### Molecular analysis

For one individual collected each in May 2023 and May 2026, nucleic acid was extracted from prosoma using DNeasy Blood & Tissue Kit (Qiagen). The barcode region of the COI gene was amplified using universal primers LCO-1490/HCO-2198 (Folmer *et al*., 1994). The polymerase chain reaction (PCR) was performed in 10 µl reactions containing 1.0 µl template DNA, 5.0 µl KOD One PCR Master Mix (TOYOBO), and 0.3 µl each primer (10 µM), using a Clontech PCR Thermal Cycler GP (TaKaRa). The thermal cycling conditions were as follows: initial step at 94–98°C (2 min); 40 cycles of denaturation at 94–98°C (10–15 s), annealing at 50–58°C (5–10 s), and extension at 68°C (10–15 s); final extension at 68°C (1 min). The PCR products were purified using ExoSAP-IT (Affymetrix), and directly sequenced by Eurofins Genomics. The obtained sequences were visually inspected using SnapGene Viewer ver. 8.2.1 (GSL Biotech LLC), and the finalized sequences were deposited in GenBank (Accession numbers: LC940353, LC940354). Chang et al. (2017) studied the population genetics of *F. albicostatus* from 16 localities in Japan, South Korea, Taiwan, and China and recovered four genetic groups. To find the phylogenetic position of Hokkaido individuals, a single sequence from each of the 16 localities (Chang et al., 2017) was downloaded from GenBank (https://blast.ncbi.nlm.nih.gov) and used to build a haplotype network. A TCS haplotype network (Templeton *et al*., 1992) was constructed using the software PopART 1.7 (Leigh *et al*., 2015). Accession numbers of the sequences used for this analysis are provided in Supplementary Table S1.

### Quantitative transect survey of barnacle abundance

To estimate the abundance of *F. albicostatus*, a line transect survey was conducted at the study site on 22 May 2026. A 129 m line transect (38.9 cm in height) on the seawall was established in the mid to upper intertidal zone (Figure 1B). A quadrat survey was subsequently conducted on 29 June 2026 to compare the abundance of *F. albicostatus* with that of co-occurring barnacle species. Ten randomly placed 10×10 cm quadrats were established. Within each quadrat, barnacle species were identified, and the number of individuals of each species was counted. *Chthamalus challengeri* and *C. dalli* were treated collectively as *Chthamalus*, and individuals smaller than about 2 mm in basal shell length were excluded from the survey as reliable species identification was not possible.

## Results

*F. albicostatus* was found during all surveys: May 2023 (*N* = 3), December 2023 (*N* = 1), May 2026 (*N* = 8), and June (*N* = 7) 2026. The shells of collected specimens were colored dark purple with white ribs (Figure 1C). The degree of rib development varied among individuals, ranging from slightly to well developed. The scutum had faint growth lines (Figure 1C). The scutal adductor ridge extended approximately halfway along the tergal margin, and was high and curved (Figure 1C). The tergum had a curved margin adjoining the carina. The apex of the spur was rounded, and the tergal depressor insertions were well developed and dentate (Figure 1C). These morphological characteristics are consistent with previous descriptions of *F. albicostatus* (Pilsbry, 1916; Utinomi 1967).

*F. albicostatus* were found sympatrically with the barnacle *Chthamalus* spp. on both boulders and seawalls (Figure 1D). *F. albicostatus* had an average abundance of 70 individuals m^-2^, together with *Chthamalus* (mean abundance 6830 individuals m^-2^) and *Balanus glandula* (220 individual m^-2^) (Figure 1E).

*F. albicostatus* is the minority species at the mid-high shores, taking up 2% abundance among all barnacle species (*Chthamalus* and *B. glandula*) (Figure 1F). The specimen collected in May 2023 (*N* = 1) had an aperture length of 0.57 cm, a basal length of 0.89 cm, and a shell height of 0.58 cm. For specimens collected in May 2026 (*N* = 6), aperture length ranged from 0.44 to 0.60 cm (mean ± SD = 0.51 ± 0.06 cm), basal diameter from 0.76 to 1.23 cm (0.88 ± 0.18 cm), and shell height from 0.31 to 0.45 cm (0.37 ± 0.06 cm). Of the eight specimens collected in May 2026, two were excluded from the measurements due to shell damage. The penis length in ethanol fixed specimens (*N* = 2, undamaged individuals only) was from 3.99 to 4.49 mm (4.24 ± 0.36 mm). All collected specimens possessed ovaries in May 2026 (*N* = 8).

COI sequences of 658 base pairs (bp) were successfully obtained from specimens collected in both 2023 and 2026. The two sequences differed by two nucleotide substitutions. To align with the available sequences of this species from previous studies, new sequences were trimmed, and the alignment had 503 bp. The new sequences from Hokkaido grouped with other sequences from Japan and Qingdao in China. The individual collected in 2023 from Hokkaido has the same haplotype with an individual from Ohtsuchi (Japan) (Figure 2).

**Fig. 2.**
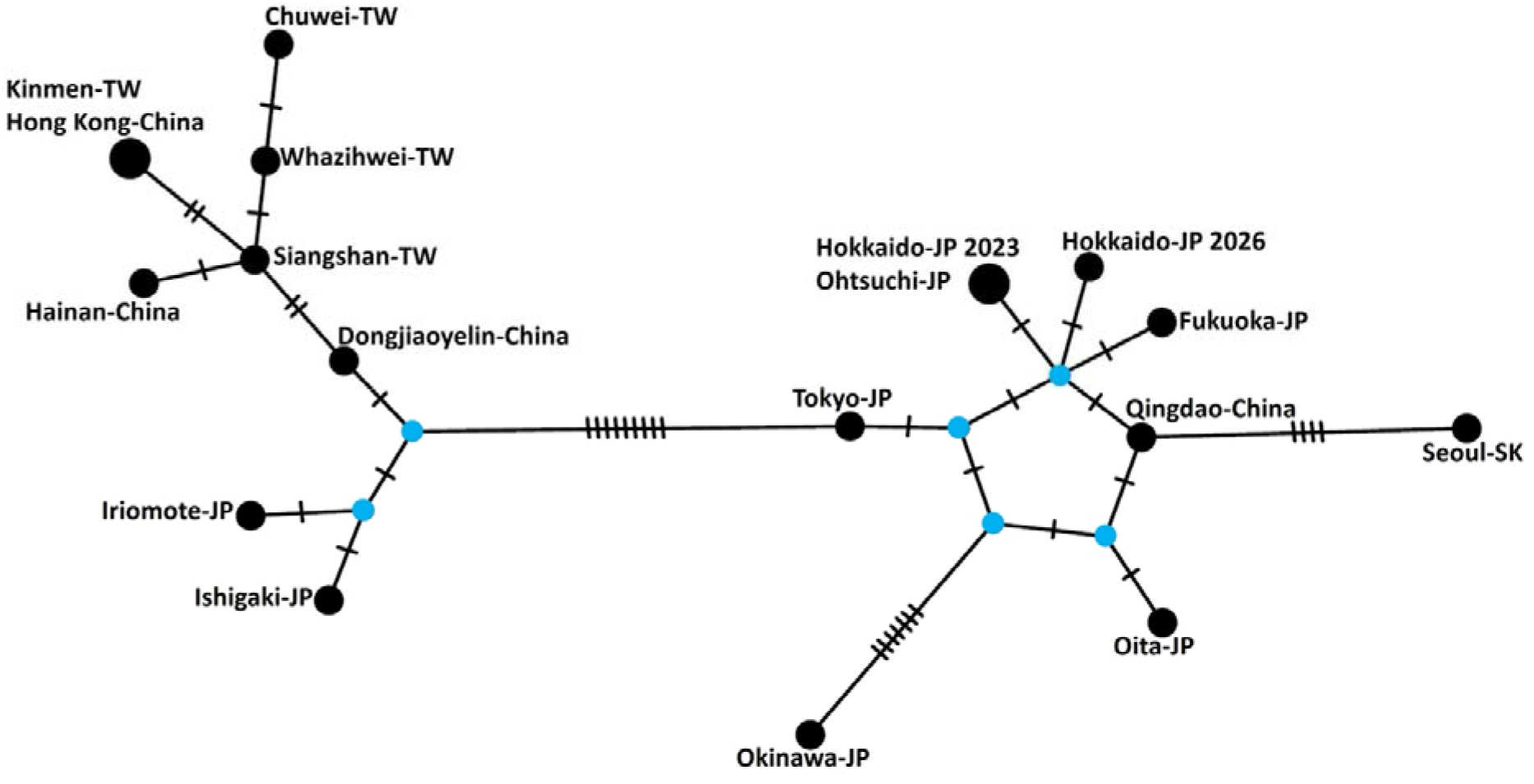
TCS haplotype network based on the COI sequences of *Fistulobalanus albicostatus* (503 base pairs), constructed using PopArt 1.7. Hatch marks indicate mutational steps; the blue circles indicate hypothetical unsampled haplotypes; the black circles, each representing a single haplotype. Abbreviations are JP: Japan; TW: Taiwan; SK: South Korea. Note: Hokkaido-JP 2023 and Hokkaido-JP 2026 are the sequences collected in the present study.

Annual trends in surface seawater and air temperatures in southern Hokkaido are shown in Figure 3 and Supplementary Table S2. For both surface seawater and air temperatures, the annual mean and maximum temperatures, and the annual number of days above the threshold temperatures increased significantly over the past 25 years (Figure 3A, B, D). The annual average air temperature increased from 9.5°C in 2000 to 11.7°C in 2025. Annual average seawater temperature increased from 10.6°C in 2000 to 11.7°C in 2025. However, minimum air and seawater temperatures did not reveal increasing trends (Figure 3C). Number of days with air temperature > 25°C increased from 2 days in 2000 to more than 20 days in 2025 (Figure 3D). Similar trends also recorded in seawater temperature that number of days that temperature > 20°C increased from 25 days in 2000 to more than 50 days in 2025. The annual number of days with air temperature below 0 °C decreased from 48 days in 2000 to fewer than 35 days in 2025 (Figure 3E). Number of days that seawater temperature below 5°C did not show clear temporal trends in the past 25 years (Figure 3E).

**Fig. 3.**
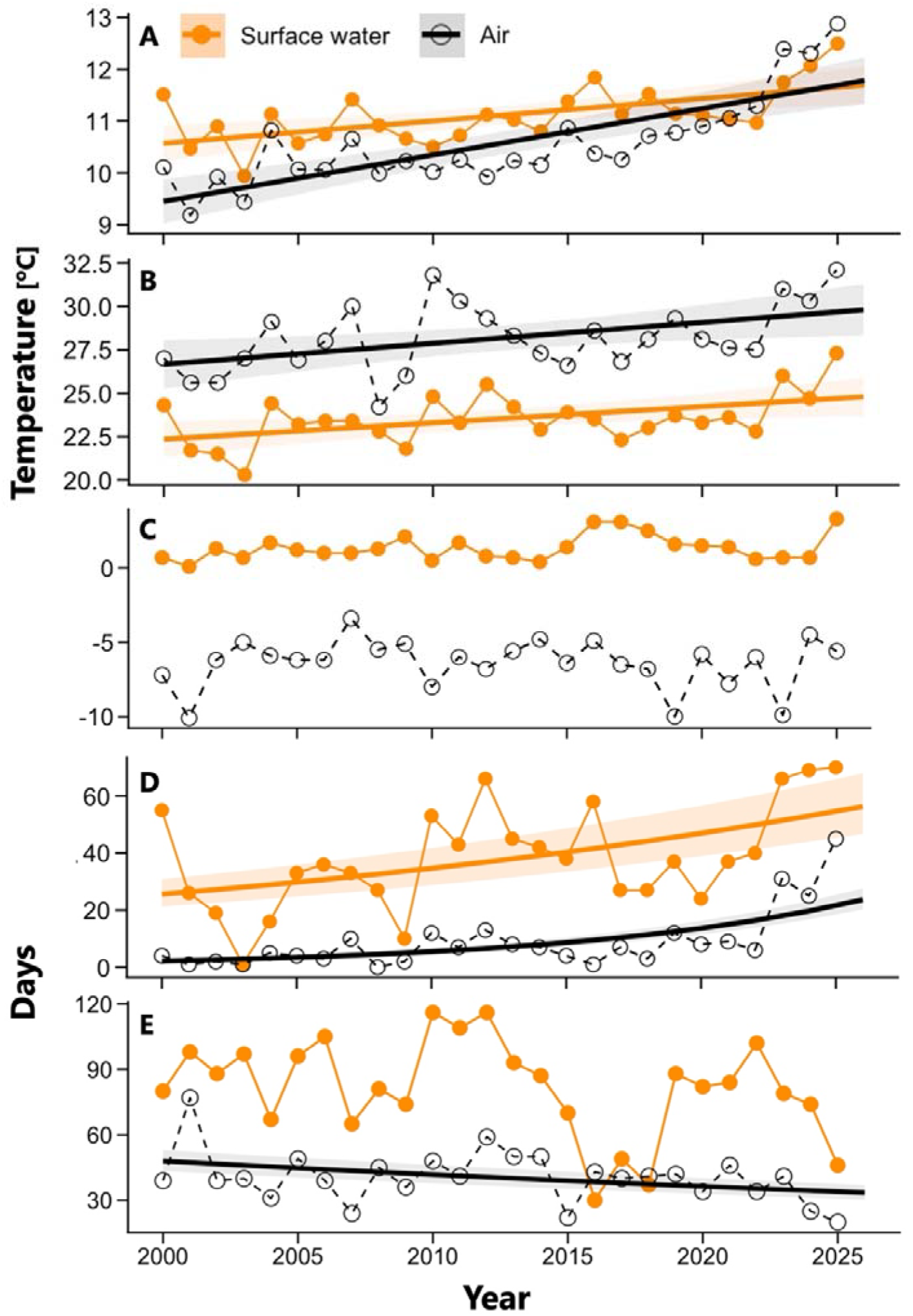
Annual trends in surface seawater and air temperatures at Usujiri, southern Hokkaido. (A) Annual mean, (B) annual maximum, (C) annual minimum temperatures, (D) annual number of days that air temperature above 20°C and seawater temperature above 25°C from 2000–2025, and (E) annual number of days below 5°C for surface water temperature and 0°C for air temperature from 2000–2025. Air temperature is shown by open black circles and dashed lines, and surface water temperature is shown by filled orange circles and solid lines. Thick lines indicate fitted model predictions from linear models (A–C), and generalized linear models with a negative binomial distribution (D–E), and shaded areas represent the 95% confidence intervals.

## Discussion

The present study identified *F. albicostatus* in Hokkaido, which is beyond its northern record (Honshu, Japan), based on both morphological characteristics and genetic information. Its occurrence in Hokkaido is unlikely to be the result of a single recruitment event, given that the species was consistently recorded at the study site between 2023 and 2026. The two samples formed two different haplotypes with close genetic affinity with individual from Ohtsuchi and Fukuoka (Honshu, Japan). This pattern supports the idea that Hokkaido individuals were likely originated from the neighboring area in Honshu, and stepwise northward range shift.

*F. albicostatus* collected in the present study had basal shell diameters ranging from 7.56 to 12.25 mm. Based on a previous study on the growth of *F. albicostatus*, the basal shell diameter increased by 6.2 mm over 11 months (Nakamura, 1997). Using this reported growth rate, *F. albicostatus* collected in Hokkaido were likely several months to more than one year post-settlement. Although this reported growth rate was obtained under relatively warm conditions (20.0–30.5°C; Nakamura, 1997), seawater temperatures in Hakodate Bay range from 4.5 to 27.3°C throughout the year (Hamao *et al*., 2022), with temperatures frequently falling below 20°C, which may result in slower growth. Therefore, the collected individuals may have been older than estimated based on the reported growth rate.

The reproductive season of *F. albicostatus* has been reported from early summer to early autumn (May–October), and individuals reach sexual maturity 17 days after settlement (Iwaki and Hattori, 1987). The specimens were collected during the early reproductive season, and possessed ovaries and well-developed penises. These observations suggest that the collected individuals were likely possessed of reproductive capability. However, *F. albicostatus* is a simultaneous hermaphrodite, and reproduces by copulation using a penis (Fraser and Chan, 2019). Given the currently low population density in Hakodate Bay, opportunities for successful mating are likely limited, although some individuals occurred in close proximity to one another (Figure 1D), potentially allowing mating. If population density increases in the future, a self-sustaining population may become established in Hakodate Bay.

One possible explanation for the occurrence of *F. albicostatus* in Hakodate Bay is the recent warming of air and seawater temperatures (Figure 3). The study site is influenced by the Tsugaru Warm Current, a warm and saline current flowing from the Sea of Japan into the North Pacific Ocean (Hamao *et al*., 2022; Figure 1A). Notably, the volume transport of this current has increased in recent years (Wakita *et al*., 2021). Several species thought to have recently expanded their distribution into Hakodate Bay via this current have been reported, including the harmful algal species *Heterosigma akashiwo* (Natsuike *et al*., 2019) and *Karenia mikimotoi* (Shimada *et al*., 2016). Similarly, larvae of *F. albicostatus* originating from populations in Honshu are likely transported to Hakodate Bay via the Tsushima and Tsugaru Warm Currents. This hypothesis is supported by haplotype analysis, which placed individuals from Hakodate Bay belonged to the same clade as those from Honshu Island.

Barnacles have a planktonic larval stage. During this stage, cyprids must locate a suitable substrate and undergo metamorphosis within a limited time window (Anderson, 1994). In *F. albicostatus*, higher water temperatures promote growth during the naupliar stage, and increase the success rate of metamorphosis to the cyprid stage (Anil *et al*., 2001; Desai *et al*., 2006). Under warmer conditions, cyprids retain their capacity to metamorphose for a longer period (Anil *et al*., 2001). Recent temperature increases may therefore have enhanced larval survival and settlement success during and after transport of *F. albicostatus* to Hakodate Bay, facilitating its establishment. Furthermore, individuals of *F. albicostatus* are exposed to air during low tide, making them vulnerable to air temperature. Although annual minimum temperatures and the annual number of days with surface water temperatures below 5°C remained unchanged (Figure 3C, E), the decrease in the annual number of days with air temperatures below 0°C may have reduced freezing stress (Figure 3E), thereby promoting survival and facilitating successful overwintering. Consequently, this reduction in subzero air temperatures in winter may have reduced these constraints and contributed to the establishment of *F. albicostatus* in Hakodate Bay.

If *F. albicostatus* has been transported via the Tsushima and Tsugaru Warm Currents, future increases in abundance and northern range expansion are likely to occur along different pathways associated with each current. After flowing northward along the coast of Honshu, the Tsugaru Warm Current passes through the Tsugaru Strait and subsequently returns to the Pacific coast of Honshu. Therefore, although larval supply to Hakodate and the Pacific coast of Honshu is likely to continue, further expansion of settlement sites along the eastern coast of Hokkaido may be limited (Figure 1A). In contrast, the Tsushima Warm Current flows northward along the Sea of Japan coast and the western coast of Hokkaido, before continuing around eastern Hokkaido and partially entering the Sea of Okhotsk as the Soya Warm Current. Consequently, if *F. albicostatus* becomes established along the Sea of Japan coast of western Hokkaido under future atmospheric and ocean warming, larval dispersal via the Tsushima and Soya Warm Currents may facilitate new population formation and further northward expansion of its distribution range. As climate change continues, warm-water species are expected to increasingly arrive and establish themselves at various locations around western and central Hokkaido. For example, central Hokkaido was also the northern limit of the stalked barnacle *Capitulum mitella* (Jones, 2000). It can be expected that the abundance or northward expansion of *C. mitella* can occur in the future.

In the present study, the occurrence of *F. albicostatus* suggests that the Tsushima Warm Current functions as a transport pathway for warm-temperate marine organisms to higher-latitude regions. The long-term trends in increasing air and seawater temperatures may be becoming increasingly possible due to recent climate change. This record provides an important baseline for assessing future changes in the coastal biota of Hokkaido.

## Data Availability Statement

The COI nucleotide sequences reported in this study have been deposited in the DDBJ/EMBL/GenBank databases under accession numbers LC940353 and LC940354. They will be publicly available upon acceptance of the manuscript.

## Acknowledgements

We sincerely thank Keiji Kobayashi for helping us with sample collection; Drs. Ryusuke Kado, Ryota Hayashi, Masashi Sekino, and Kohei Matsuno for valuable comments and suggestions; and the Usujiri Fisheries Station at Hokkaido University, especially Atuya Miyajima, for providing the temperature data; the Uehiro Laboratory for Oceanography at Hokkaido University for the support. We are also grateful to the members of the Plankton, and the Humans and the Ocean Laboratories at Hokkaido University, especially Dr. Shingo Akita and Yuta Jozawa, for allowing us to use their facilities and for their kind guidance.

## Author contributions

MT conceived this study, carried out the fieldwork, conducted laboratory measurements, generated COI sequence data, and drafted the initial manuscript. AS conducted haplotype analysis. BKK plotted the graphs of the abundance. AS and BKK contributed to the writing and revision of the manuscript. All authors approved the final manuscript.

## Ethical standards

The experiments were conducted by following the National University Corporation Hokkaido University Regulations on Animal Experimentation.

## References

Anderson DT (1993) Barnacles: structure, function, development and evolution. London: Chapman & Hall.

Anil AC, Desai D and Khandeparker L (2001) Larval development and metamorphosis in Balanus amphitrite Darwin (Cirripedia; Thoracica): significance of food concentration, temperature and nucleic acids. Journal of Experimental Marine Biology and Ecology 263, 125–141.

Chan BKK and Leung PT (2007) Antennular morphology of the cypris larvae of the mangrove barnacle Fistulobalanus albicostatus (Cirripedia: Thoracica: Balanomorpha). Journal of the Marine Biological Association of the United Kingdom 87, 913–915.

Chan BKK, Tsang LM and Chu KH (2007) Morphological and genetic differentiation of the acorn barnacle Tetraclita squamosa (Crustacea, Cirripedia) in East Asia and description of a new species of Tetraclita. Zoologica Scripta 36, 79–91.

Chang YW, Chan JS, Hayashi R, Shuto T, Tsang LM, Chu KH and Chan BKK (2017) Genetic differentiation of the soft shore barnacle Fistulobalanus albicostatus (Cirripedia: Thoracica: Balanomorpha) in the West Pacific. Marine Ecology 38, e12422.

Chen IC, Hill JK, Ohlemüller R, Roy DB and Thomas CD (2011) Rapid range shifts of species associated with high levels of climate warming. Science 33, 1024–1026.

Desai D, Khandeparker L and Shirayama Y (2006) Larval development and metamorphosis of Balanus albicostatus (Cirripedia: Thoracica); implications of temperature, food concentration and energetics. Journal of the Marine Biological Association of the United Kingdom 86, 335–343.

Fraser CM and Chan BKK (2019) Too hot for sex: mating behaviour and fitness in the intertidal barnacle Fistulobalanus albicostatus under extreme heat stress. Marine Ecology Progress Series 610, 99–108.

Fredston□ Hermann A, Selden, R, Pinsky M, Gaines SD and Halpern BS (2020) Cold range edges of marine fishes track climate change better than warm edges. Global Change Biology 26, 2908–2922.

Figueira WF and Booth DJ (2010) Increasing ocean temperatures allow tropical fishes to survive overwinter in temperate waters. Global Change Biology 16, 506–516.

Foster BA and Willan RC (1979). Foreign barnacles transported to New Zealand on an oil platform. New Zealand journal of marine and freshwater research, 13, 143–149.

Hadley W (2016) ggplot2: Elegant Graphics for Data Analysis. New York: Springer-Verlag.

Hamao Y, Morimoto K, Tatamisashi S, Wakita M, Kasai A and Matsuno K (2024) Seasonal changes in the protist communities of Hakodate Bay, southern Hokkaido, from 2020 to 2022. Regional Studies in Marine Science 78, 103775.

Hayashi R (2017) First documentation of the barnacle Fistulobalanus albicostatus (Cirripedia: Balanomorpha) as an epibiont of loggerhead sea turtle Caretta caretta. Marine Biodiversity 47, 157–158.

Henry DP and McLaughlin PA (1975) The barnacles of the Balanus amphitrite complex (Cirripedia, Thoracica). Zoologische verhandelingen 141, 1–254.

Iwaki T and Hattori H (1987) First maturity and initial growth of some common species of barnacles in Japan. Bulletin of the Faculty of Fisheries, Mie University 14, 11–19.

Jones DS, Hewitt MA and Sampey A (2001) A checklist of the Cirripedia of the South China Sea. Raffles Bulletin of Zoology 48, 233–308.

Kado R (2003) Invasion of Japanese shores by the NE Pacific barnacle Balanus glandula and its ecological and biogeographical impact. Marine ecology progress series 249, 199–206.

Kotwicki L, Weslawski JM, Włodarska-Kowalczuk M, Mazurkiewicz M, Wenne R, Zbawicka M, Minchin D and Olenin S (2021) The re-appearance of the Mytilus spp. complex in Svalbard, Arctic, during the Holocene: The case for an arrival by anthropogenic flotsam. Global and Planetary Change 202, 103502.

Leigh JW, Bryant D and Nakagawa S (2015) POPART: Full-Feature Software for Haplotype Network Construction. Methods in Ecology and Evolution 6, 1110–1116.

Lenoir J, Bertrand R, Comte L, Bourgeaud L, Hattab T, Murienne J and Grenouillet G (2020) Species better track climate warming in the oceans than on land. Nature ecology & evolution 4, 1044–1059.

McGillycuddy M, Popovic G, Bolker BM and Warton DI (2025) Parsimoniously fitting large multivariate random effects in glmmTMB. Journal of Statistical Software 112, 1–19.

Brooks ME, Kristensen K, van Benthem KJ, Magnusson A, Berg CW, Nielsen A, Skaug HJ, Maechler M and Bolker BM (2017) glmmTMB Balances Speed and Flexibility Among Packages for Zero-inflated Generalized Linear Mixed Modeling. The R Journal 9, 378–400.

Natsuike M, Kanamori M and Shimada H (2019) Red tide and seasonal occurrence of the harmful raphidophyte Heterosigma akashiwo in Hakodate Bay, Hokkaido. Scientific Reports of Hokkaido Fisheries Research Institutes (Japan) 95, 11–17.

Pilsbry HA (1916) The barnacles (Cirripedia): contained in the collections of the US National museum; including a monograph of the American species. Bulletin of the United States National Museum 93, 1–366.

Pinsky ML, Selden RL and Kitchel ZJ (2020) Climate-driven shifts in marine species ranges: scaling from organisms to communities. Annual review of marine science 12, 153–179.

R Core Team (2024) R: A Language and Environment for Statistical Computing. Vienna, Austria: R Foundation for Statistical Computing. Available at https://www.R-project.org/ (accessed online 1 July 2026).

Rantanen M, Karpechko AY, Lipponen A, Nordling K, Hyvärinen O, Ruosteenoja K, Vihma T and Laaksonen A (2022) The Arctic has warmed nearly four times faster than the globe since 1979. Commun Earth Environ 3, 168.

Shimada H, Kanamori M, Yoshida H and Imai I (2016) First record of red tide due to the harmful dinoflagellate Karenia mikimotoi in Hakodate Bay, southern Hokkaido, in autumn 2015. Nippon Suisan Gakkaishi 82, 934–938.

Sorte CJ, Williams SL and Carlton JT (2010) Marine range shifts and species introductions: comparative spread rates and community impacts. Global Ecology and Biogeography 19, 303–316.

Templeton AR, Crandall KA and Sing CF (1992) A cladistic analysis of phenotypic associations with haplotypes inferred from restriction endonuclease mapping and DNA sequence data. III. cladogram estimation. Genetics 132: 619–633.

Tøttrup AP, Chan BKK, Koskinen H and Høeg JT (2010) ‘Flying barnacles’: implications for the spread of non-indigenous species. Biofouling 26, 577–582.

Utinomi H (1967) Comments on some new and already known cirripeds with emended taxa, with special reference to the parietal structure. Publication of Seto Marine Biological Laboratory 15, 199–237.

Wakita M, Sasaki K, Nagano A, Abe H, Tanaka T, Nagano K, Sugie K, Kaneko H, Kimoto K, Okunishi T, Takada M, Yoshino J and Watanabe S (2021) Rapid reduction of pH and CaCO3 saturation state in the Tsugaru Strait by the intensified Tsugaru Warm Current during 2012–2019. Geophysical Research Letters 48, e2020GL091332.

