## Supplemental Table S1, and S2 for "Northward expansion of the barnacle *Fistulobalanus albicostatus* in Japan"

**Supplementally materials**

Tables

| **Sampling locality** | | **GenBank accession number** |
| --- | --- | --- |
| Japan | Hakodate | LC940353 |
|  |  | LC940354 |
|  | Ohtsuchi | KX405194.1 |
|  | Tokyo | KX405357.1 |
|  | Fukuoka | KX405305.1 |
|  | Oita | KX405270.1 |
|  | Okinawa | KX405321.1 |
|  | Iriomote | KX405397.1 |
|  | Ishigaki | KX405210.1 |
| South Korea | Seoul | JX503003.1 |
| Taiwan | Siangshan | KX405463.1 |
|  | Wazihwei | KX405484.1 |
|  | Chuwei | KX405403.1 |
|  | Kinmen | KX405363.1 |
| China | Qingdao | JQ035512.1 |
|  | Hong Kong | KX405193.1 |
|  | Hainan | KX405366.1 |
|  | Dongjiaoyelin | KX405369.1 |

**Table S1.** Sampling localities and GenBank accession numbers of COI gene sequences used to build a haplotype network.

| **Target** | **Factor** | **Estimate** | **SE** | ***t* or *Z*** | ***p*** |
| --- | --- | --- | --- | --- | --- |
| **Annual mean temperature** | | | | | |
| Surface seawater | Intercept | –75.726 | 23.230 | –3.260 | 0.003 |
|  | Year | 0.043 | 0.012 | 3.738 | 0.001 |
| Air | Intercept | –169.782 | 28.477 | –5.962 | <0.001 |
|  | Year | 0.090 | 0.014 | 6.333 | <0.001 |
| **Annual maximum temperature** | | | | | |
| Surface seawater | Intercept | –164.861 | 68.557 | –2.405 | 0.024 |
|  | Year | 0.094 | 0.034 | 2.748 | 0.011 |
| Air | Intercept | –214.432 | 93.253 | –2.299 | 0.030 |
|  | Year | 0.121 | 0.046 | 2.602 | 0.016 |
| **Annual minimum temperature** | | | | | |
| Surface seawater | Intercept | –75.628 | 43.565 | –1.736 | 0.095 |
|  | Year | 0.038 | 0.022 | 1.767 | 0.090 |
| Air | Intercept | 18.239 | 88.828 | 0.205 | 0.839 |
|  | Year | –0.012 | 0.044 | –0.277 | 0.784 |
| **Annual number of days above the threshold temperatures** | | | | | |
| Surface seawater  (>20 ºC) | Intercept | –57.445 | 25.750 | –2.231 | 0.026 |
|  | Year | 0.030 | 0.013 | 2.372 | 0.018 |
| Air  (>25 ºC) | Intercept | –182.494 | 36.729 | –4.969 | <0.001 |
|  | Year | 0.092 | 0.018 | 5.025 | <0.001 |
| **Annual number of days below the threshold temperatures** | | | | | |
| Surface seawater  (<5 ºC) | Intercept | 29.117 | 15.125 | 1.925 | 0.054 |
|  | Year | –0.012 | 0.008 | –1.635 | 0.102 |
| Air  (<0 ºC) | Intercept | 31.552 | 14.164 | 2.228 | 0.026 |
|  | Year | –0.014 | 0.007 | –1.966 | 0.049 |

**Table S2.** Results of linear models (LMs) and generalized linear models (GLMs) examining temporal trends (year effects) in environmental conditions at Usujiri, southern Hokkaido, Japan. LMs were applied to temperature variables (annual mean, maximum, and minimum temperatures), and GLMs were applied to count variables (annual number of days above and below the threshold temperatures), assuming a negative binomial distribution with a log link function.
